# Unique transcriptional changes in coagulation cascade genes in SARS-CoV-2-infected lung epithelial cells: A potential factor in COVID-19 coagulopathies

**DOI:** 10.1101/2020.07.06.182972

**Authors:** Ethan S. FitzGerald, Amanda M. Jamieson

**Affiliations:** Division of Biology and Medicine, Department of Molecular Microbiology and Immunology, Brown University, Providence, Rhode Island, United States

## Abstract

Severe acute respiratory syndrome coronavirus 2 (SARS-CoV-2) has rapidly become a global pandemic. In addition to the acute pulmonary symptoms of COVID-19 (the disease associated with SARS-CoV-2 infection), pulmonary and distal coagulopathies have caused morbidity and mortality in many patients. Currently, the molecular pathogenesis underlying COVID-19 associated coagulopathies are unknown. While there are many theories for the cause of this pathology, including hyper inflammation and excess tissue damage, the cellular and molecular underpinnings are not yet clear. By analyzing transcriptomic data sets from experimental and clinical research teams, we determined that changes in the gene expression of genes important in the extrinsic coagulation cascade in the lung epithelium may be important triggers for COVID-19 coagulopathy. This regulation of the extrinsic blood coagulation cascade is not seen with influenza A virus (IAV)-infected NHBEs suggesting that the lung epithelial derived coagulopathies are specific to SARS-Cov-2 infection. This study is the first to identify potential lung epithelial cell derived factors contributing to COVID-19 associated coagulopathy.

**GRAPHICAL ABSTRACT:** 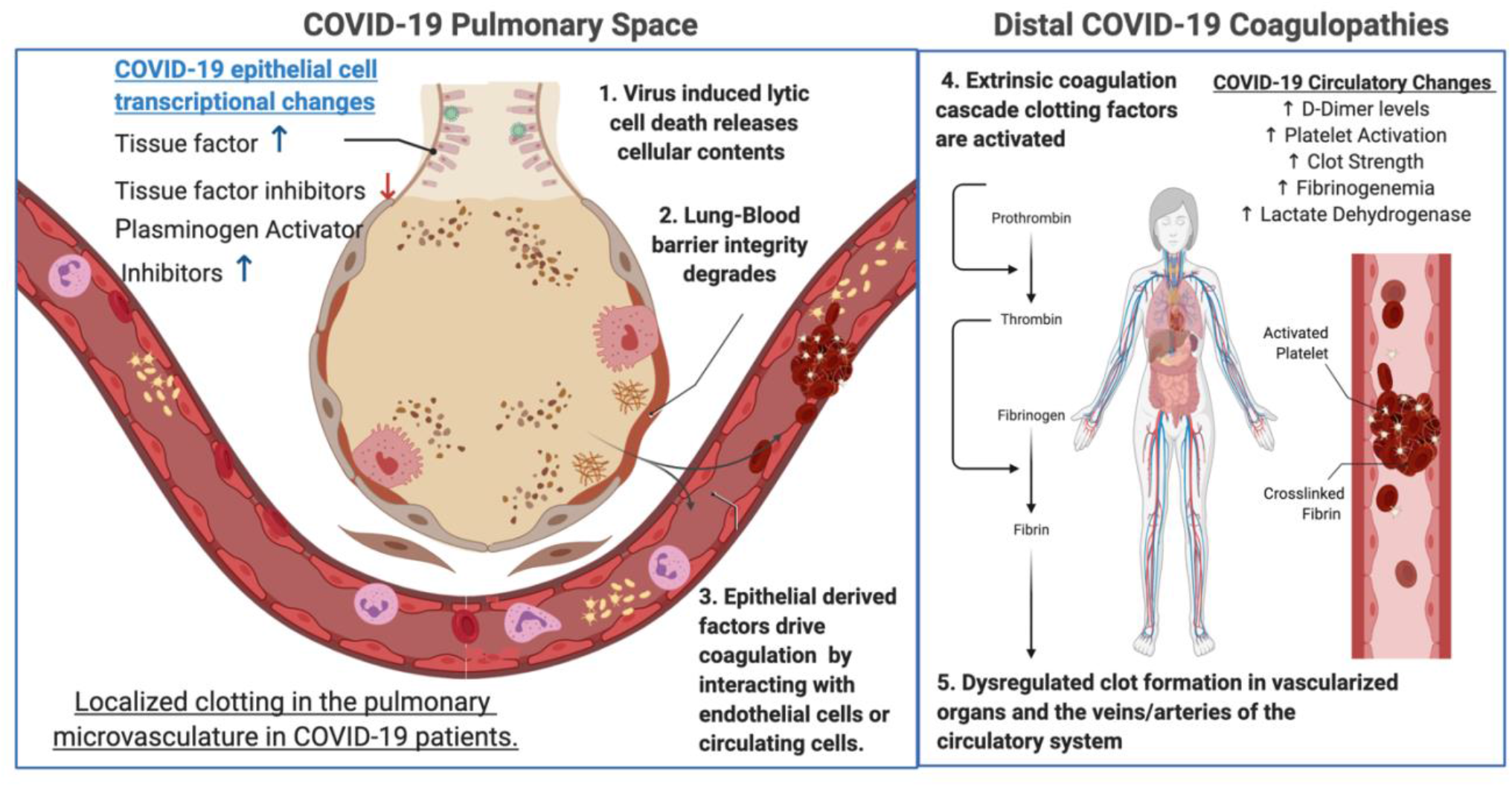

**AUTHOR SUMMARY:** *Why was this study done?:*

- Severe acute respiratory syndrome coronavirus 2 (SARS-CoV-2) has rapidly become a global pandemic.
- In addition to the acute pulmonary symptoms of COVID-19 (the disease associated with SARS-CoV-2 infection), pulmonary and distal coagulopathies have caused morbidity and mortality in many patients.
- Currently, the molecular pathogenesis underlying COVID-19 associated coagulopathies are unknown. Understanding the molecular basis of dysregulated blood coagulation during SARS-CoV-2 infection may help promote new therapeutic strategies to mitigate these complications in COVID-19 patients.

*What did the researchers do and find?:*

- We analyzed three publicly available RNA sequencing datasets to identify possible molecular etiologies of COVID-19 associated coagulopathies. These data sets include sequencing libraries from clinically isolated samples of bronchoalveolar lavage fluid (BALF) and peripheral blood mononuclear cells (PBMCs) from SARS-CoV-2 positive patients and healthy controls. We also analyzed a publicly available RNA sequencing dataset derived from *in vitro* SARS-CoV-2 infected primary normal human bronchial epithelial (NHBE) cells and mock infected samples.
- Pathway analysis of both NHBE and BALF differential gene expression gene sets. We found that SARS-CoV-2 infection induces the activation of the extrinsic blood coagulation cascade and suppression of the plasminogen activation system in both NHBEs and cells isolated from the BALF. PBMCs did not differentially express genes regulating blood coagulation.
- Comparison with influenza A virus (IAV)-infected NHBEs revealed that the regulation of the extrinsic blood coagulation cascade is unique to SARS-CoV-2, and not seen with IAV infection.

*What do these findings mean?:*

- The hyper-activation of the extrinsic blood coagulation cascade and the suppression of the plasminogen activation system in SARS-CoV-2 infected epithelial cells may drive diverse coagulopathies in the lung and distal organ systems.
- The gene transcription pattern in SARS-CoV-2 infected epithelial cells is distinct from IAV infected epithelial cells with regards to the regulation of blood coagulation.

## INTRODUCTION

In December of 2019, a novel respiratory coronavirus, designated SARS-CoV-2, emerged in Wuhan China.^1^ It has since spread globally causing major societal shutdowns and >10 million confirmed infections with >500,000 recorded deaths.^2,3^ Initial clinical reports described the symptomology of COVID-19 (the disease caused by SARS-CoV-2) as a pneumonia presenting with fever, fatigue, shortness of breath, and a dry cough.^4^ Severe cases are often complicated by acute respiratory distress syndrome (ARDS) and cytokine storm associated hyper-inflammation, with many patients requiring mechanical ventilation and ICU admission due to hypoxia and pneumonia.^4,5^ The pathology of COVID-19 also impacts organ systems and tissues beyond the lung, including the kidneys, gut, liver, and brain.^6–9^ Many of the most concerning distal pathologies associated with SARS-CoV-2 infection have been associated with increased blood coagulation and clotting. These diverse coagulopathies have included venous, arterial, and microvascular thromboses of idiopathic origin.^8,10–14^

Blood coagulation is primarily regulated by three highly interconnected molecular signaling pathways, platelet activation, the coagulation cascade, and fibrinolysis.^15,16^ Many excellent review articles on the molecular players in this process are available.^17–25^ The extrinsic blood coagulation pathway is effected through a cascading activation of zymogen coagulation factors, which is balanced by endogenously encoded zymogen inhibitors. The end result of the extrinsic coagulation cascade is the formation of crosslinked fibrin clots mediated by activated thrombin. Plasmin suppresses blood coagulation and clotting via proteolytic degradation of these cross linked fibrin blood clots. Increases in pro-coagulant biomarkers are known to be associated with greater risk of mortality for patients suffering acute lung injury (ALI).^26–28^ Modulation of blood coagulation and fibrinolysis have previously been proposed as therapeutic strategies for the treatment of ALI.^29^ Many research teams and medical associations have recommended elevated D-dimer and other serum markers of coagulation be measured as biomarkers of COVID-19 disease severity for in-patient testing. Blood thinning treatments such as heparin have begun to be administered prophylactically to minimize the risk of COVID-19 associated coagulopathies, and clinical trials are ongoing to investigate the efficacy of common blood thinning medications and anti-coagulants at mitigating COVID-19 morbidity and mortality.^30–33^ (ClinicalTrials.gov Identifiers: NCT04333407 & NCT04365309)

Coagulopathies concomitant to hyper-inflammatory injury such as ARDS and sepsis have been hypothesized to synergize due to interactions of inflammation and the extrinsic coagulation cascade.^34,35^ It has been broadly theorized that COVID-19 associated coagulopathies are indirectly induced by the acute inflammation and pulmonary tissue damage associated with SARS-CoV-2, but precise mechanisms underlying this severe COVID-19 disease phenotype have remained elusive.^36,37^ The identification of the tissue or cellular origins of the signal transducing molecules that drive dysregulated blood coagulation will be critical to understanding the pathogenesis of SARS-CoV-2-induced coagulopathies. To this end, we have performed post-hoc analysis on publicly available transcriptomics datasets of SARS-CoV-2-infected normal human bronchial epithelial cells (NHBEs), COVID-19 patient bronchoalveolar lavage fluid (BALF) and COVID-19 peripheral blood mononuclear cells (PBMCs), with the goal of generating hypotheses regarding the possible etiology of SARS-CoV-2 induced coagulopathies.^38,39^ We found that there is a clear transcriptional signature of dysregulated blood coagulation cascade signaling in NHBEs that are infected with SARS-CoV-2. These cells are infected *in vitro*, so this gene signature is not influenced by interactions with immune cells. However, transcriptional analysis of BALF cells revealed many similarities in the regulation of genes important in the coagulation cascade. In contrast, PBMCs isolated from blood do not show this gene signature, indicating that the coagulopathy defect is derived from lung signals. In addition, comparison with transcriptional data from NHBE cells infected with influenza A virus (IAV) revealed that the dysregulation of genes important in coagulation in lung epithelial cells is not generalizable to all respiratory infections.^39^ Our study demonstrates that changes to the lung epithelium directly caused by SARS-CoV-2 infection may be responsible for the coagulopathy seen in COVID-19 patients.

## METHODS

### Xiong et al. – RNA-seq analysis of BALF and PBMCs from SARS-CoV2 infected patients

BALF and PBMC sequencing data were generated through the purification of cells and subsequent RNA-sequencing libraries from SARS-CoV-2 infected patients in the Zhongnan Hospital of Wuhan University as described in Xiong *et al.*^38^ These analyses were performed on samples collected as part of standard treatment and diagnostic regimens. No extra burden was imposed on patients.

Briefly, PBMC cells were purified from peripheral blood samples obtained from 3 patients and 3 healthy donors via Ficoll density gradient centrifugation. Purified PBMC in the buffy coat were then transferred to a falcon tube and washed with PBS before RNA purification using Trizol and Trizol LS reagents according to the manufacturer’s protocol. BALF cells were isolated from patients by injecting 2% lidocaine solution into the right middle lobe or left lingular segment of the lung for local anesthesia. 100ml of room temperature sterile saline was used to lavage the right middle lobe or left lingular segment of the lung before transfer to sterile containers. Three BALF control samples isolated from healthy volunteers were downloaded from a publicly available NCBI dataset at sample accession SRR10571724, SRR10571730, and SRR10571732.^40^

Library preparation for BALF and PBMC samples were performed manually using 1μg of total RNA as input. The library prep involved poly-A enrichment using oligo-dt capture probes, heat fragmentation, first and second strand synthesis with adapter ligation, and PCR library amplification. After library size selection, the prepared dsDNA libraries were denatured and circularized to allow for rolling circle amplification to form DNA nano-balls (DNBs). The DNBs were then quantified and sequenced on MGISEQ-200 platforms. PBMC and BALF samples were sequenced on either MGISEQ-2000 or Illumina NovaSeq platforms. Raw sequencing data were submitted to the Chinese Academy of Science’s Genome Sequence Archive (GSA) (COVID+ BALF - GSA Accession CRP001417; PBMCs – GSA Accession CRA002390).

### Blanco-Melo et al. - RNA-seq analysis of SARS-CoV2 infected NHBEs cultured in-vitro

Normal human bronchial epithelial (NHBE) cells isolated from a 78 year old Caucasian woman were cultured under non-differentiating conditions in bronchial epithelial growth media supplemented with BEGM SingleQuots. (Lonza, CC-4175). SARS-CoV-2 isolate USA-WA1/2020 (NR-52281) was propagated in Vero E6 cells, and viral titers were determined via plaque assay on Vero E6 cells. NHBE (5×10^5^) cells were infected with SARS-CoV-2 at a multiplicity of infection of 2 for 24 hours and or mock infected in their culture media. Total RNA was then isolated via TRIzol extraction and Direct-zol RNA Miniprep Kit (Zymo research) cleanup.

Library preparation for NHBE samples was performed using the TruSeq Stranded mRNA Library Prep Kit (Illumina) with Poly-A enrichment, according to the manufacturer’s instructions. Libraries were sequenced on the Illumina NextSeq 500 platform. Raw sequencing data were submitted to the National Center for Biotechnology Information’s Gene Expression Omnibus (GEO Accession - GSE63473).

### Blanco-Melo et al. - RNA-seq analysis of H1N1 infected NHBEs cultured in vitro

Normal human bronchial epithelial (NHBE) cells isolated from a 78 year old Caucasian woman were cultured under non-differentiating conditions in bronchial epithelial growth media supplemented with BEGM SingleQuots. (Lonza, CC-4175). A/Puerto Rico/8/1934 (PR8) influenza virus infection was performed on cells in their culture media at a multiplicity of infection of 3 for 12 hours. As described for SARS-CoV-2 infection, Total RNA was then isolated via TRIzol extraction and Direct-zol RNA Miniprep Kit (Zymo research) cleanup.

Library preparation for NHBE samples infected with influenza A virus (IAV) was performed using the TruSeq Stranded mRNA Library Prep Kit (Illumina) with Poly-A enrichment, according to the manufacturer’s instructions. Libraries were sequenced on the Illumina NextSeq 500 platform. Raw sequencing data were submitted to the National Center for Biotechnology Information’s Gene Expression Omnibus (GEO Accession - GSE63473).

### Transcriptional analysis pipeline

The analysis pipeline described below was used to analyze HBEC, PBMC, and BALF data sets for functional enrichments in differentially expressed genes. Sequencing reads were downloaded from the National Center for Biotechnology Information’s Sequence Read Archive or the Chinese Genomic Data Sharing Initiative’s Genome Sequence Archive in their raw fastq format. Read adapter and quality trimming was performed using the Trim Galore! package^41^ and Sequencing quality was assessed using the FASTQC and MULTIQC packages.^42,43^ All read files were deemed to be of sufficient quality for analysis to proceed. Sequence alignment to the GRCh38 reference transcriptome was performed using the Salmon read alignment software and differential gene expression analysis was performed using DESeq2 and the tximport packages.^44,45^ For all differential expression analyses, the infected group of each cell type was compared to the uninfected group of the same cell type.

PantherDB was used to perform functional enrichment analysis using the Biological Processes Gene Ontology annotation set, with a user supplied gene list of all differentially expressed genes with an adjusted P value of less than then 0.2.^46,47^ The adjusted P value cut-off of .2 was selected to include genes that may exhibit biologically significant changes in gene expression in the Gene Ontology analysis. Individual gene plots and heat maps were generated in R using the pheatmap or ggplot2 R packages to directly evaluate each gene’s differential expression.^48,49^

## RESULTS

### Coagulation pathway gene expression in human bronchial epithelial cells is impacted by infection with SARS-CoV-2

In order to determine the impact that SARS-CoV-2 infection has on key factors in the coagulation cascade we examined the transcription of genes that are important in the regulation of hemostasis and venous thrombosis, including the extrinsic coagulation pathway and the plasminogen activation system. Supplemental Table 1 lists the functions of all the genes within the blood coagulation cascade that are differentially expressed in SARS-CoV-2 infected NHBEs relative to mock-infected controls. These genes are part of the regulation of blood coagulation gene ontology (GO) term (GO:0030193), which was identified as significantly enriched by PantherDB functional enrichment analysis of all NHBE differentially expressed genes (P.adj > 0.2) (Figure 1A). Importantly, the transposition of gene expression directionality onto pathway maps for the extrinsic blood coagulation cascade and plasminogen activation pathway (Figure 1B and Supplemental Table 1), illustrate how infected respiratory epithelial cells may drive this coagulopathy in COVID-19 infection. Most notably, tissue factor is significantly transcriptionally upregulated while balancing inhibitory proteins are either unmodified or significantly downregulated by epithelial cells during infection. Additionally, while plasminogen activating proteins are significantly upregulated, plasminogen activating inhibitors and localizing receptors are also transcriptionally increased. The combination of these transcriptional modifications during SARS-CoV-2 infection may significantly contribute to coagulopathies associated with COVID-19.

**Figure 1:**
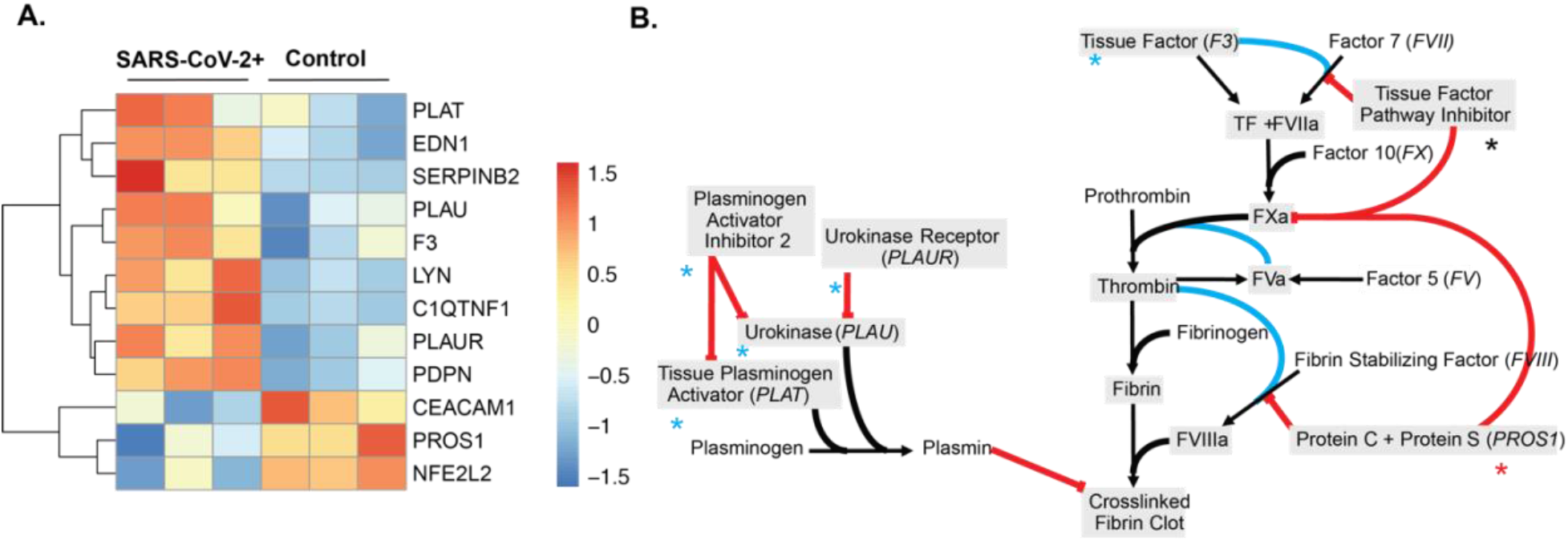
The gene expression profile of differentially expressed genes within the enriched the regulation of blood coagulation GO term for SARS-CoV-2 infected NHBE cells. (A) Heatmap of all differentially expressed genes (Padj > .2) enriched in the regulation of blood coagulation GO term. False discovery rate was calculated in PantherDB using functional enrichment analyzing all Biological Process GO Terms. (B) Pathway map of the extrinsic blood coagulation cascade (right) and the plasminogen activation system (left) with overlaid expression values. Blue asterisks indicate upregulation, black asterisks indicate no change, and red asterisks indicate down regulation.

### Regulation of tissue factor in NHBEs infected with SARS-CoV-2

One clear factor that could be impacting the coagulation cascade is the increased expression of *F3* in SARS-CoV-2 infected NHBEs (Figure 2A). The *F3* gene encodes the Tissue Factor protein, which is secreted by a variety of tissue cells to initiate the extrinsic coagulation cascade. This signaling cascade is carefully regulated by the balance of tissue factor protein with several endogenously encoded inhibitor proteins that suppress the signaling cascade. The first inhibitory protein in this cascade is the *TFPI* gene, which encodes the Tissue Factor Pathway Inhibitor (TFPI) protein. TFPI acts to suppress blood coagulation by inhibiting the activation of factor VII at the head of the signaling cascade. *TFPI* transcription is not significantly different in COVID infected epithelial cells relative to mock infected cells (Figure 2B). The maintenance of homeostasis between tissue factor and TFPI is essential for the maintenance of vascular systems without excessive clotting, and the observed increase of tissue factor without corollary increases of TFPI could significantly contribute to the induction of clotting in COVID-19 patients.^50^

**Figure 2:**
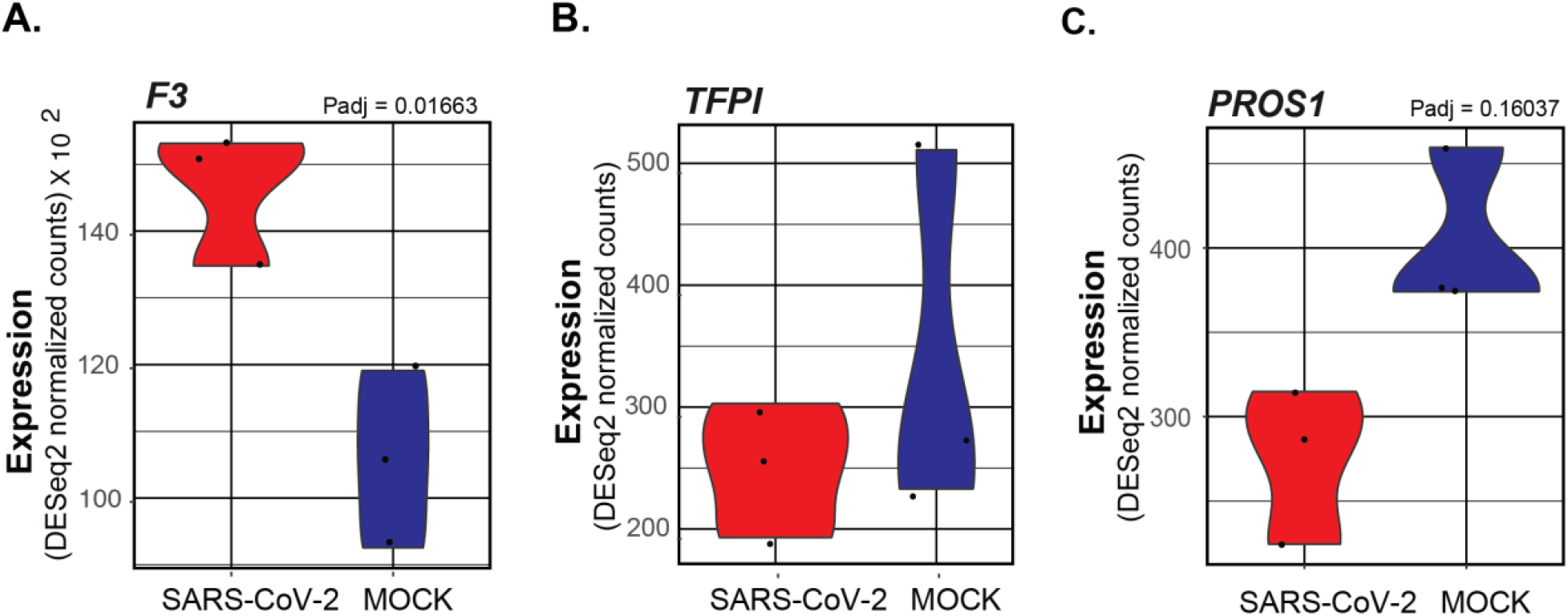
Violin plots depicting raw counts of reads mapping to key regulators of the extrinsic blood coagulation cascade in mock infected and SARS-CoV-2 infected NHBE cells. Raw counts were normalized to library size in the DESeq2 software package. Adjusted P values for all differentially expressed genes were also calculated within DESeq2. Genes lacking P values are not differentially expressed. Images were generated using GGPlot2 in the R studio environment.

### Decreased expression of PROS1 in NHBEs infected with SARS-CoV-2

Another critical suppressor of the extrinsic blood coagulation cascade, the *PROS1* gene which encodes Protein S, was also found to be downregulated in SARS-CoV-2 infected NHBEs (Figure 2C). Protein S is a vitamin K dependent glycoprotein with homology to Factors VII, IX, and X in the coagulation cascade. Its primary function is to antagonize the coagulation cascade by complexing with Protein C. The complex, known as Activated Protein C, acts to inhibit the maturation of pro-coagulation factors Va and VIIIa. This results in the suppression of both pro-thrombin maturation and thrombin activity. However, the activity of both protein C and protein S is required for this effect.^18^ It also is known to promote the activity of TFPI.^23^ Interestingly, the binding of protein S also contributes to efferocytic clearance of apoptotic cells by mediating membrane dynamics between macrophages and epithelial cells. Its activity is highly anti-inflammatory in this capacity, and decreased expression of *PROS* expression may further exacerbate COVID-19 related pathology through diverse mechanisms.^51^

### NHBE cell regulation of plasminogen by SARS-CoV-2 infection

Additionally, within the Regulation of Blood Coagulation GO Term, several genes regulating the activity of plasminogen were identified. Both *PLAU* (encoding Urokinase) and *PLAT* (encoding the tissue plasminogen activator protein) are observed to be significantly increased in SARS-CoV-2 NHBEs (Figures 3A and 3B). NHBEs infected with SARS-CoV-2 significantly upregulate expression of *SERPINB2*, which encodes the protein Plasminogen Activator Inhibitor 2 (PAI-2) (Figure 3C). PAI-2, is known to be a potent inhibitor of both Urokinase and tissue plasminogen activator, acting through the proteolytic inactivation of plasminogen activators. (DOI: 10.1007/s00018-004-4230-9) While PAI-2 is most commonly localized within the cytoplasm, the increased release of markers of membrane permeable cell death such as lactate dehydrogenase may provide evidence for the increased secretion of other cytoplasmic proteins such as PAI-2. (doi:10.1001/jama.2020.1585) The expression of *PLAUR*, a receptor localizing activated urokinase to the cell membrane, is also significantly increased in SARS-CoV-2 infected NHBEs (Figure 3D). The localized activity of PAI-2 may significantly inhibit the effect of PLAU/PLAUR in complex and thereby contribute to the formation of pulmonary embolisms and distal coagulopathies.

**Figure 3:**
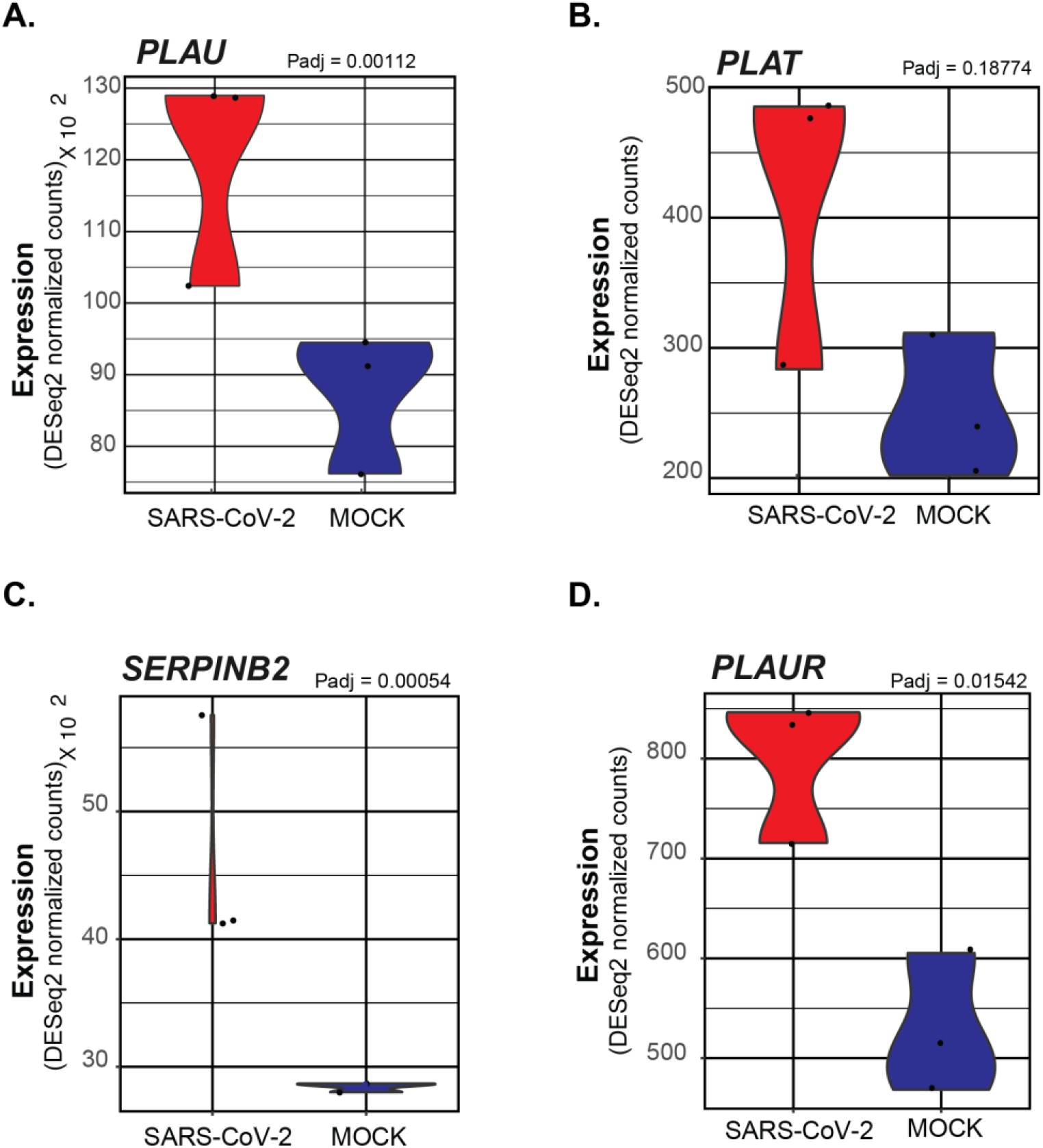
Violin plots depicting raw counts of reads mapping to regulators of the plasminogen activation system in mock infected and SARS-CoV-2 infected NHBE cells. Raw counts were normalized to library size in the DESeq2 software package. Adjusted P values displayed for significant differences were also calculated within DESeq2. Images were generated using GGPlot2 in the R studio environment.

### Regulation of blood coagulation by cells isolated from the BALF of COVID-19 Patients

Plotting of the subset of genes in the Regulation of Blood Coagulation (GO:0030193) GO term that were initially found to differentially expressed in SARS-CoV-2 infected NHBEs, revealed a clear pattern of transcriptional regulation in cells isolated via BALF of COVID-19 patients as well (Figure 4A). In addition, plotting of the expression data of all BALF differentially expressed genes (P.adj > .2) in the Regulation of Blood Coagulation GO term revealed clear regulation of the blood coagulation cascade occurring locally within the bronchoalveolar space (Figure 4B). Many of these expression signatures recapitulate the findings we observed when analyzing *in vitro* SARS-CoV-2 infected NHBE cells.

**Figure 4:**
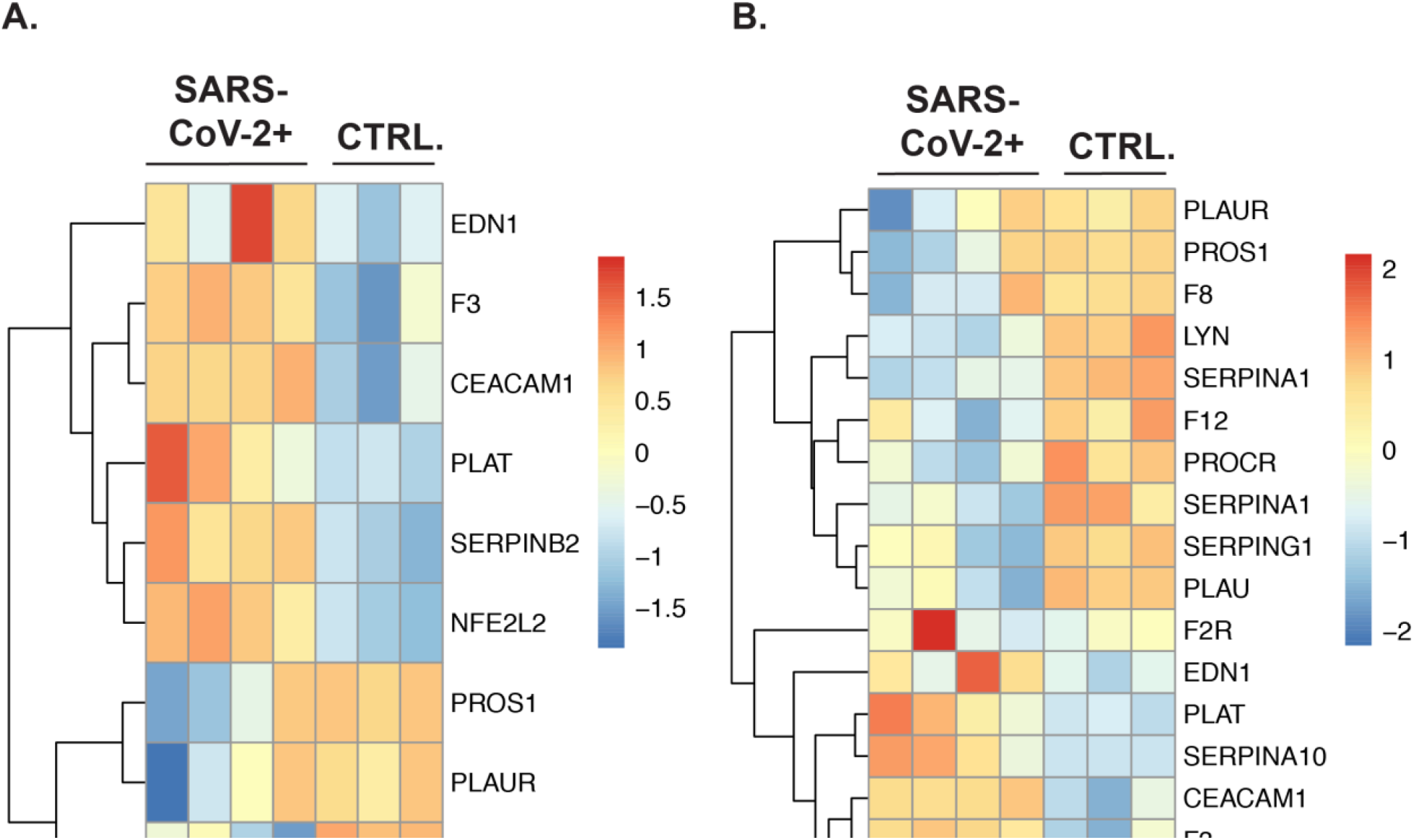
The gene expression profile of differentially enriched genes from RNA isolated from the BALF of COVID-19 patients. (A) The genes included in this heatmap were identified as enriched in the regulation of blood coagulation GO term for NHBE cells infected with SARS-CoV-2. The expression data presented in the heat map demonstrates the expression profile of these genes in BALF derived samples. (B) The genes presented in this heatmap represent all BALF differentially expressed genes (P. adj > .2) that are also included in the Blood Coagulation GO Term (GO:0007596). PantherDB functional enrichment analysis of BALF differentially expressed genes (P. adj > .2) including all Biological Process GO terms did not identify the Blood Coagulation GO term as statistically enriched.

These include the upregulation of pro-coagulation genes such as *F3* (tissue factor) and *SERPINB2*, along with the downregulation of inhibitory genes such as *PROS1* and *PLAUR*, and *PLAT*. Additionally, the gene *PROCR*, a receptor that augments the inhibitory activity of protein S and protein C, was found to be suppressed in the BALF during infection. Unlike the NHBE data set there is increased expression of *TFPI* and *PLAT* in the BALF from SARS-CoV-2, indicating that hosts are actively signaling to suppress coagulation during COVID-19. However, the increasing appearance of coagulopathies in COVID-19 patients indicate that this signaling is often insufficient to prevent morbidity and mortality.

In analyzing these data, is important to note that BALF samples contain a complex mixture of resident and recruited immune cells, along with damaged tissue cells that have been freed from the membrane, often concomitantly with cell death processes. Prior research indicates that during SARS-CoV-1 infection, there is significant epithelial denudation which may result in a greater fraction of cells in BALF samples from infected individuals containing epithelial cells and type 2 pneumocytes.^52,53^ As such, the transcriptional signature in analyzed BALF samples represents a bulk averaging of this complex mixture. Additionally, given the impact of varying co-morbidities when examining patient derived samples, (including smoking status, age, and non-related pre-existing conditions) further analysis of additional BALF patient samples is required to confirm these observations.

### Analysis of coagulation pathway gene expression in PBMCs

In order to determine if coagulation pathway gene expression was changed in circulating immune cells we analyzed sequencing datasets generated from COVID-19 patient purified PBMCs as described in Xiong et al. PantherDB Functional enrichment analysis found no significant enrichment of genes regulating or effecting blood coagulation in PBMC datasets in PBMCs from COVID-19 patients vs. controls (Figure 5; BP Full In supplement). The primary publication associated with these datasets describe expected induction of genes relating to the hyper-inflammatory response associated with ARDS and the induction of regulated cell death in immune cells. From these data, we concluded it is unlikely that circulating immune cells during SARS-CoV-2 infection are inducing blood coagulation through the secretion of signals activating the extrinsic or intrinsic blood coagulation cascade in response to systemic inflammation.

**Figure 5:**
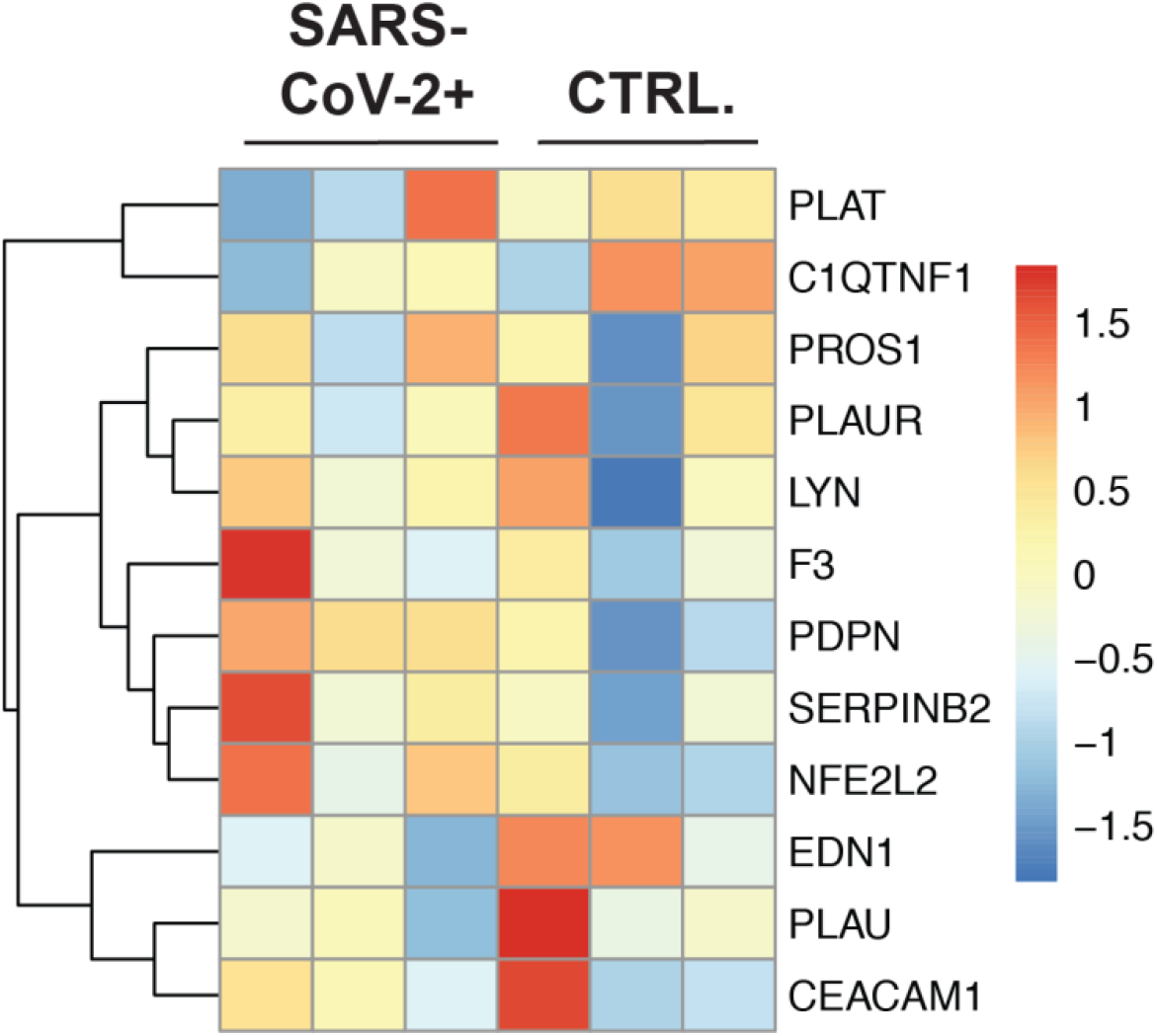
The gene expression profile of differentially enriched genes from RNA isolated from PBMCs of COVID-19 patients. **(A)** The genes included in this heatmap were identified as enriched in the regulation of blood coagulation GO term for NHBE cells infected with SARS-CoV-2. The expression data presented in the heat map demonstrates the expression profile of these genes in PBMC derived samples.

### Infection of human lung epithelial cells with influenza A virus does not impact coagulation pathway gene expression

In order to determine if these transcriptional changes are specific for SARS-CoV-2 or are more generalizable to respiratory viruses that infect the lung epithelium we analyzed data sets from IAV infected NHBE cells. PantherDB Functional enrichment analysis found no significant enrichment of genes regulating or effecting blood coagulation in these sequencing datasets when comparing IAV infected and uninfected NHBEs (Full functional enrichment results in supplement). Furthermore, heatmap plotting of genes found to be differentially expressed in NHBE cells during SARS-CoV-2 infection (Figure 1A), did not reveal any notable patterns or differential expression signatures in the context of IAV infection (Figure 6). These findings are consistent with the lack of severe coagulopathies associated with IAV infection in the clinic and further support the notion that the induction of coagulopathies during SARS-CoV-2 infection is independent of systemic inflammation common to both infections. This indicates that COVID-19 associated coagulopathies may be triggered by changes in lung epithelial cell transcription patterns uniquely induced by SARS-CoV-2 infection.

**Figure 6:**
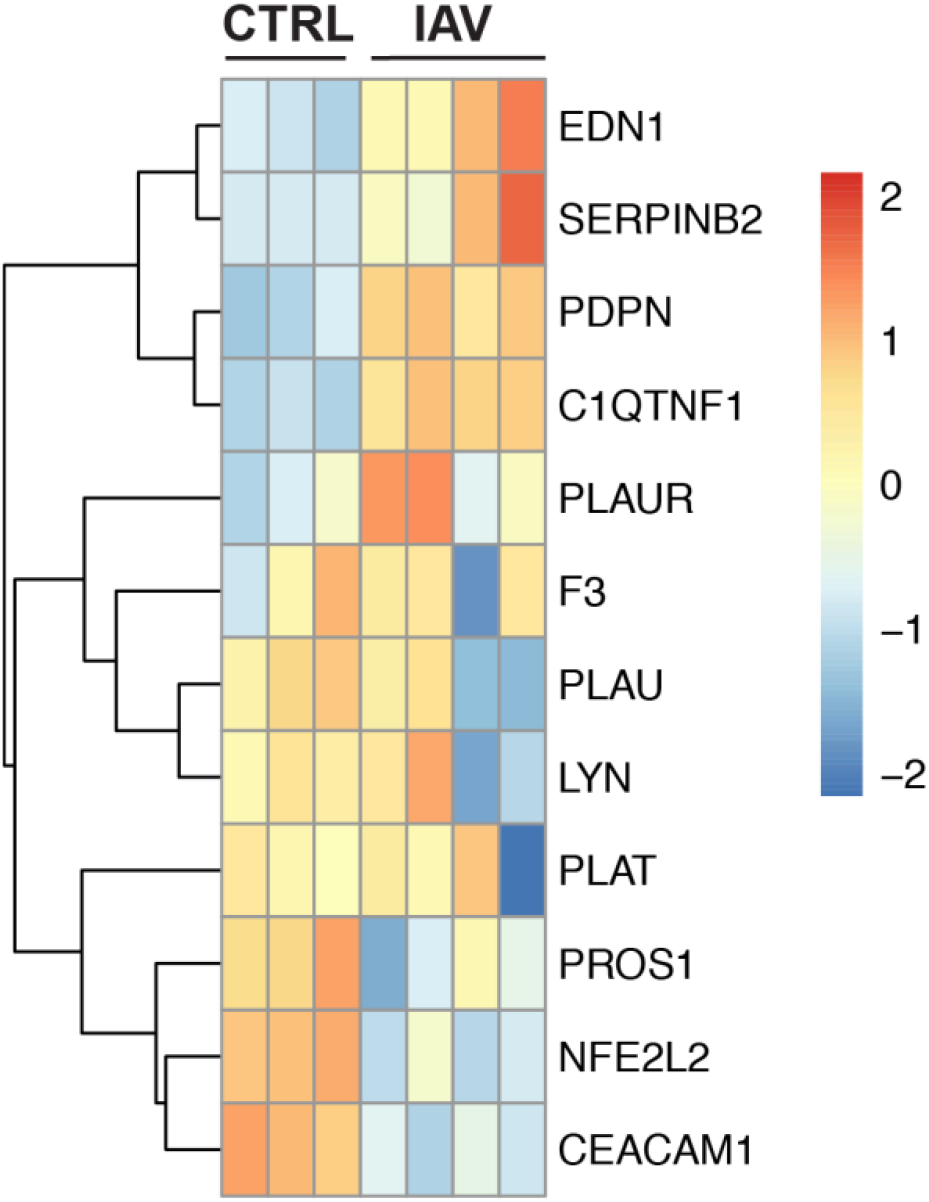
The gene expression profile of differentially enriched genes from RNA isolated from NHBE cells infected with PR8 IAV. (A) The genes included in this heatmap were identified as enriched in the regulation of blood coagulation GO term for NHBE cells infected with SARS-CoV-2. The expression data presented in the heat map demonstrates the expression profile of these genes in NHBE cell cultures that are mock infected or infected with PR8 IAV at a multiplicity of infection of 3.

## DISCUSSION

The data presented here demonstrate that SARS-CoV-2 infection of human bronchial epithelial cells may drive three key molecular responses promoting coagulopathies associated with COVID-19; (1) induction of the extrinsic coagulation cascade through the activation of tissue factor signaling without compensatory tissue factor pathway inhibitor expression; (2) the suppression of anticoagulation signaling through the down regulation of Protein S in the pulmonary space; and (3) the upregulation of plasminogen inactivation proteins and localization factors. Such activities by infected pulmonary epithelial cells *in vivo* could significantly predispose patients to the hyper-coagulation that has been associated with SARS-CoV-2. In addition, our data indicate that the same genes are not upregulated by human bronchial epithelial cells infected with IAV, indicating that the severity of coagulopathy in COVID-19 patients may be derived from changes in infected lung epithelial cells.

There is mounting evidence that coagulation defects are a significant and severe pathology of COVID-19. An early clinical correspondence published in the New England Journal of Medicine reported that 5 New York City patients under the age of 50 presented with large vessel arterial stroke from March 23 to April 7, 2020.^10^ Since then, reports of COVID-19 associated coagulopathies in the young and old have proliferated globally, with reports describing acute pulmonary embolism in the microvasculature of the lung, as well as cerebral, renal, and bowel localized embolic disease.^8,11,12^ Reports have shown that acute pulmonary thromboembolism presents in 30% of severe clinical COVID-19 patients by pulmonary CT angiography. These emboli were found to be associated with elevation of serum D-dimer, which is produced during the degradation of crosslinked fibrin clots by enzymes such as plasmin.^13^ Some preliminary reports have also found that biomarkers of coagulation such as clot strength, platelet and fibrinogen contributions to clots, and elevated d-dimer levels are significantly increased with ARDS caused by COVID-19.^14^ A diverse spectrum of proinflammatory mediators shown to be dramatically upregulated in COVID-19 and other coronavirus pathologies are also known to contribute to tissue factor induced hypercoagulability.^54^ Many other molecular factors increased with SARS-CoV-2 infection, including phosphatidylserine exposure, interferon expression, ICAM expression, angiotensin II expression, and complement activation, are also known to “decrypt” tissue factor from its inactive form on the surface of tissue cells.^18^ Such “coagulation-inflammation-thrombosis” circuit feedback loops coupled with the multiple zymogen activation mediated feedback loops within the extrinsic blood coagulation cascade, could significantly contribute to the induction of COVID-19 coagulopathy in patients.^18^ However, further investigation of the activation of the extrinsic blood coagulation cascade or the inhibition of plasmin by respiratory epithelial cells is required to validate these hypotheses.

To best treat patients it is necessary to understand the tissue, cellular, and molecular underpinnings of COVID-19 pathology. In order to investigate the role that systemic and lung cells play in coagulopathy we analyzed three distinct data sets. The first dataset, published in Xiong *et al.* performed transcriptome sequencing on peripheral blood mononuclear cells (PBMCs) and bronchoalveolar lavage fluid (BALF) cells isolated from human patients infected with SARS-CoV-2.^38^ The second data set, published in Blanco-Melo et al. performed transcriptome sequencing on commercially purchased normal human bronchial epithelial cells isolated from a 78 year old woman infected with SARS-CoV-2 and IAV.^39^

Coagulation cascade induction is thought to be necessary during ARDS or ALI, and may be protective.^55^ However, when it becomes dysregulated it can be damaging. Also, systemic coagulation defects can cause severe pathologies. For instance, tissue factor and other genes within the extrinsic coagulation cascade and fibrinolysis pathway were previously found to contribute to ALI in a murine model of coronavirus infection.^59^ ARDS is often associated with increased biomarkers of coagulation and fibrinolysis. For instance, increases in pro-coagulant biomarkers are known to be associated with greater risk of mortality for patients suffering acute lung injury. Pulmonary edema fluids and plasma from patients with acute lung injury have also been shown to contain lesser amounts of anti-coagulant protein C and higher amounts of plasminogen activator inhibitors, likely secreted from epithelial and endothelial pulmonary cells.^26–28^ However, our comparison of gene signatures from IAV infected NHBEs with those from SARS-CoV-2 infected NHBEs indicate that SARS-CoV-2 may be unique from other respiratory viruses in terms of the risk of coagulation defects. Changes to the lung epithelium, separate from inflammatory immune responses, may increase the risk of coagulopathies.

The role of the lung epithelium in coagulation defects has not been fully explored, however several lines of evidence demonstrate that it may play a key role in some instances. Lung epithelial cell lines have been shown to have increased expression of TF after incubation with pulmonary edema fluid from ARDS patients.^56^ In addition, mouse models demonstrate that lung epithelial-derived TF may play an important role in tissue protection during ALI caused by LPS.^57^ In vitro experiments with human epithelial cells indicate that TF may also be important for lung epithelial basal cell survival.^58^ Taken together these lines of evidence suggest that while induction of the extrinsic coagulation cascade by lung epithelial cells may be an important host response during some stages of infection, but SARS-CoV-2 can cause such profound changes that this leads to hyper-coagulation and systemic pathologies.

Other important players in regulation of the coagulation cascade are vascular endothelial cells. Endothelial cells are known to release soluble tissue factor in response to cytokines, which may further contribute to extrinsic coagulation cascade induced coagulopathies in COVID-19 patients.^59^ The possibility of direct endothelial cell infection by SARS-CoV-2, which has been shown to occur *in vitro* and may occur *in vivo*, should also be considered as a possible mechanism for the induction of hyper-coagulation signals driving COVID-19 associated coagulopathies.^60^ Indirect damage of endothelial cells during acute lung injury associated with ARDS could also drive these signals. However, to our knowledge, there are currently no RNA-sequencing datasets with infected endothelial cell cultures or tissue available. Such data sets would be invaluable in determining how endothelial cells respond to epithelial cell coagulation signals or how endothelial cells directly modulate hyper-coagulation associated with SARS-CoV-2 infection.

Many researchers have proposed that lytic regulated cell death by respiratory epithelial cells, particularly pyroptosis, may play a significant role in COVID-19 pathogenesis.^61^ During lytic cell death many intracellular pathogen associated molecular patterns (PAMPs) and damage associated molecular patterns (DAMPs) typically isolated within cell membranes are released. However, it is also underappreciated that diverse intracellular and membrane bound contents are also released in addition to PAMPs and DAMPs. It is possible that proteins such as tissue factor, plasminogen activating inhibitors, and pro-coagulant factors may be released into the pulmonary space during COVID-19 induced lytic cell death of epithelial cells. If this is the case, such factors may drive paracrine signaling to nearby endothelial cells. This could further exacerbate coagulation systemically by inducing the secretion of activated coagulation cascade zymogens and thrombin into the blood. Such factors could also enter the blood stream directly near damaged endothelial tissues in the lung. Epithelial cell derived hyper-coagulation factors and plasminogen inhibitors also may drive local pulmonary hyper-coagulation and further exacerbate tissue destruction in the lung during SARS-CoV-2 infection.

Further investigation of pulmonary endothelial, epithelial, and immune cell responses to SARS-CoV-2 will be essential for unraveling the mystery of COVID-19 induced hyper-coagulation. The current BALF data is from bulk sequencing, which can obscure cell-type specific gene signatures. In addition, further characterization of SARS-CoV-2 infected human respiratory epithelial cells in physiologically relevant *in vitro* systems, such as air liquid interface cultures, would increase our understanding of the interaction of SARS-CoV-2 with the lung epithelium. Culture of vascular endothelial cells could be used to determine the role that either direct infection of endothelial cells or exposure to inflammatory cues may play in causing coagulopathies. Such approaches would facilitate sequencing, imaging, or proteomic investigations of the activity of tissue factor, plasminogen activation inhibitor 2, and plasminogen activators in infected lung cells.

While further investigation is required to determine if epithelial cell signaling are driving coagulopathy, several possible clinical appoaches could be considered after further validation. For instance, serum samples from SARS-CoV-2 positive patients with severe COVID-19 symptoms or coagulopathies could be assayed to determine the concentration and activation state of zymogens effecting the extrinsic coagulation cascade.^62^ These approaches could similarly be utilized to design blood panels for stratifying COVID-19 in-patient coagulopathy risk or monitoring of embolic disease risk. Identifying the molecular and cellular factors that drive SARS-CoV-2 induced coagulopathy is essential, both from the perspective of understanding the biology behind SARS-CoV-2 and in terms of clinical treatments. The data in this study demonstrate that SARS-CoV-2 has a unique impact on pulmonary cells. Furthering our understanding of SARS-CoV-2 pathology, and the central role that the lung epithelium may play on this pathology will be essential in determining the ideal treatment regimens for COVID-19.

## Supporting information

HBEC Blood Coag DEG Annotation Table

BALF-PantherDB BP Complete - Padj <.2

HBEC_IAV-PantherDB BP Complete - Padj <.2

PBMC-PantherDB BP Complete - Padj <.2

Covid_DEGs_BALF

Covid_DEGs_NHBE

Covid_DEGs_PBMC

