## Supplemental Table 1 for "Unique transcriptional changes in coagulation cascade genes in SARS-CoV-2-infected lung epithelial cells: A potential factor in COVID-19 coagulopathies"

| Gene Symbol - Protein Product | Functional Description |
|---|---|
| **PLAT - Tissue Plasminogen activator** | Processes plasminogen into plasmin, which is a serine protease that helps to break down clots in the blood. Recombinant PLAT is used as a anti-clotting therapy. (doi: 10.1080/08998280.2011.11928729) |
| **EDN1 - Endothelin 1** | Vasoconstrictor that is proteolytically processed to be effected. Known contributor to pulmonary vasculature constriction which has been treated with a small molecule (Bosentan). (doi: 10.1186/rr44 ; doi: 10.1183/23120541.00060-2018) |
| **SerpinB2 - Plasminogen activator inhibitor-2** | A coagulation factor which works by inhibiting PLAT's serine protease function and inbiting urokinase. Hyperactivity is known to contribute to thrombosis through this inhibition. (DOI: 10.1055/s-0031-1276589) |
| **PLAU - Urokinase** | Serine protease which participates in the maturation of plasmin to help break down blood clots. It also activates proteases that can degrade ECM and may promote wounding responses in COVID+ settings. Inhibitors have been developed to inhibit PLAUs contribution to invasive cancer. Urokinase has also been applied to suppress clotting and thrombosis. (https://doi.org/10.3389/fonc.2018.00024 ; doi: 10.3892/mmr.2018.8414 ; doi: 10.3389/fneur.2017.00371) |
| **F3 - Tissue Factor** | Master regulator of the Extrinsic blood coagulation cascade via its secretion which activates a protease signaling cascade to promote blood coagulation. (htt |
| **LYN - Tyrosine-protein kinase Lyn** | Src Kinase known to interact with platelets during their activation. (doi: 10.1074/jbc.M109.098756) |
| **C1QTNF1 - Complement 1q and TNF Related 1** | Increased serum levels have been identified as a marker of high risk for major cardiovascular events in diabetic patients. This includes clotting, stroke, and thrombosis. (DOI:https://doi.org/10.1016/j.atherosclerosis.2019.06.377) |
| **PLAUR - Urokinase receptor** | Membrane receptor which binds to Urokinase (PLAU) and restricts its activity to the localized microenvironment. This activity promotes local tissue and ECM degradation during wound healing. PLAUR and PLAU in complex can be proteolytically cleaved to release active suPAR into circulation to degrade clots. (DOI: 10.1183/13993003.01571-2015 , https://doi.org/10.1182/blood.V92.6.2075) |
| **PDPN - Protoplanin** | Mucin like protein. Interaction of PDPN with platelet expressed CLEC2 is known to induce platelet activation activation and drives blood coagulation in vitro and in vivo. Observed to promote cancer related thrombosis. Also known to contribute to the localization of ECM degrading components such as urokinase in invadopodia. (https://doi.org/10.1038/s41467-017-02402-6 ; doi: 10.1182/blood.2019001388 ; doi: 10.1038/onc.2014.388) |
| **CEACAM1 - Carcinoembryonic antigen-related cell adhesion molecule 1** | Cytoplasmic membrane glyco-protein that contributes to the inhibition of immune receptor tyrosine-based activation motif signaling and limit thrombus growth by suppressing platelet activation and adhesion through receptor mediated interactions. (doi: 10.1182/blood-2008-06-165043) |
| **PROS1 - Protein S** | Vitamin K dependent glycoprotein that has homology with Factor VII, IX, and X in the coagulation cascade. Its primary function is to contribute to the anti-coagulation pathway in its free form, where it complexes with Protein C to inhibit the maturation of pro-coagulation factors Va and VIIIa . The binding of protein S also contributes to efferocytic clearance of apoptotic cells by mediating membrane dynamics between macrophages and epithelial cells. Its activity is highly anti-inflammatory in this capacity. (doi: 10.1055/s-0037-1604092) |
| **NFE2L2 - NRF2** | Transcription factor regulating the expression of antioxidants. Generally drives many cytoprotective genes and is known to help promote the suppression of blood coagulation through AGE/RAGE signaling and the suppression of stress responses relating to oxidation. (doi: 10.3390/ijms20133208) |

https://vivo.brown.edu/display/ajamieso

**Supplemental Table 1**
